# Cumulative microscopy reveals cellular states in fibroblasts from patients with genetic disorders

**DOI:** 10.1101/2025.08.04.668398

**Authors:** Karen Akopyan, Susann Karlberg, Svetlana Vakkilainen, Fulya Taylan, Minna Pekkinen, Outi Mäkitie, Arne Lindqvist

**Affiliations:** Department of Cell and Molecular Biology, Karolinska Institutet, Stockholm, Sweden; Children’s Hospital, Pediatric Research Center, University of Helsinki and Helsinki University Hospital, Helsinki, Finland; Research Program for Clinical and Molecular Metabolism, Faculty of Medicine, University of Helsinki, Helsinki, Finland; Folkhälsan Research Center, Institute of Genetics, Helsinki, Finland; Department of Molecular Medicine and Surgery, Karolinska Institutet, and Department of Clinical Genetics and Genomics, Karolinska University Hospital, Stockholm, Sweden

## Abstract

Analysis of cellular states and signaling trajectories can provide insights into causes of disease. We developed cumulative microscopy, a method to perform cyclical imaging without elution or quenching steps. Cumulative microscopy computationally extracts individual signals from accumulating fluorescence during sequential imaging. We used cumulative microscopy to quantitatively assess cell cycle and stress markers in individual primary fibroblasts from patients with rare genetic proliferative disorders with increased cancer risk. Neural network-based analysis of cumulative microscopy data suggested that cells from patients with Cartilage-hair hypoplasia (CHH), but not Mulibrey Nanism (MUL), showed replication stress. We analyzed cell states and cell trajectories and found that a subset of cells from patients with CHH showed spontaneous replication stress, followed by cell cycle exit in both G1 and G2 phase. We note that replication stress potentially could underlie both proliferative defects and increased cancer risk in CHH patients and conclude that cumulative microscopy is an efficient, quantitative, and generalizable approach to multiplex microscopy.

**Graphical abstract:** 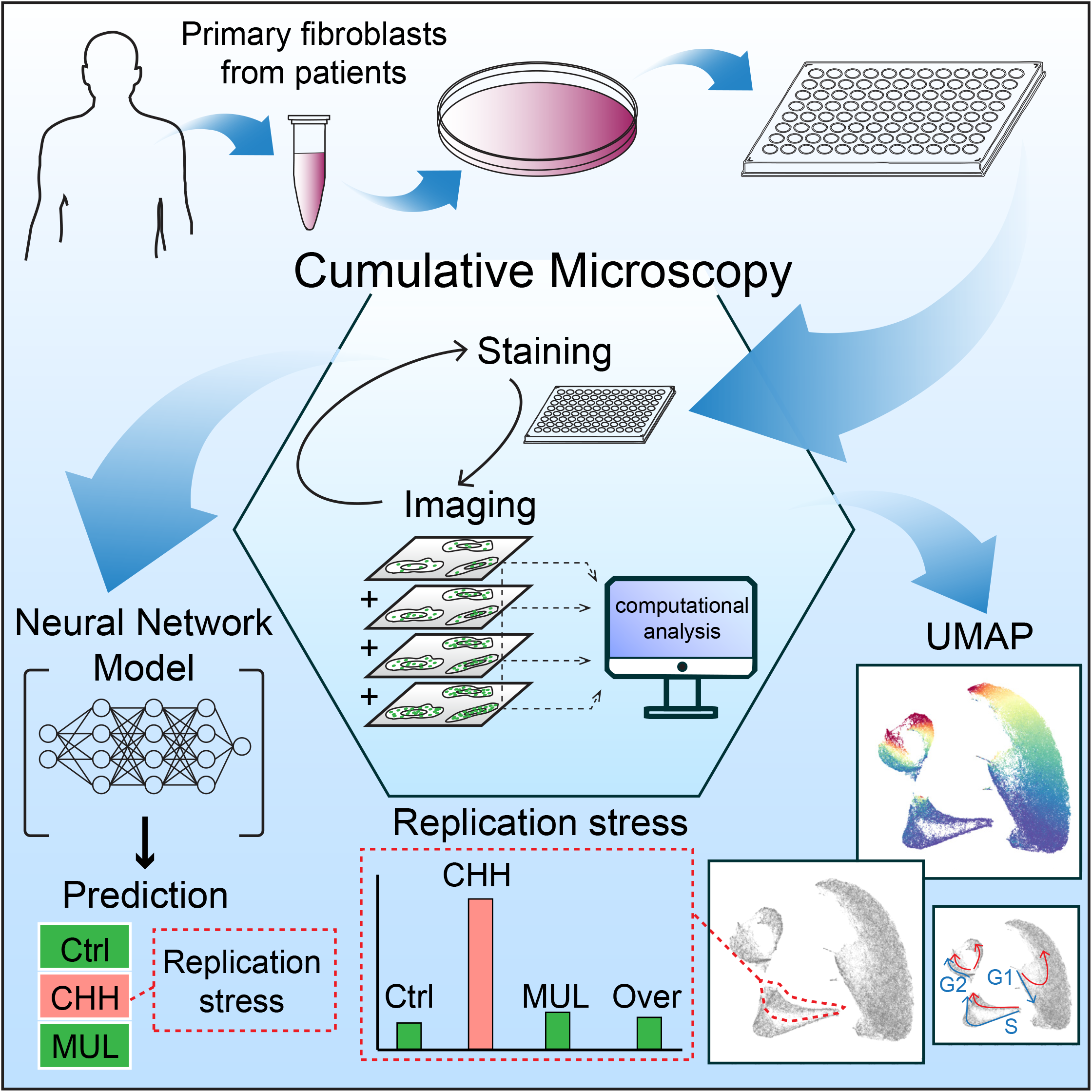

## Introduction

The cell forms the basic unit of life. The collective action of cells impacts on a wide range of processes in health and disease. To understand and model cellular behavior, a large number of different parameters can be described that collectively outline various cellular states (Rafelski and Theriot, 2024). Transitions between cellular states can be used to model eventual cell fates, while also accounting for various responses in a heterogenous population of cells (Rafelski and Theriot, 2024; Movasat et al., 2025).

Several thousand rare genetic disorders have been identified but understanding of the mechanistic connection between genotype and phenotype remains often incomplete (Haendel et al., 2020). These include cartilage-hair hypoplasia (CHH; OMIM #250250) and Mulibrey Nanism (MUL; OMIM #253250). Whereas CHH is caused by mutations in the non-coding RNA *RMRP* that forms part of a large RNA processing complex (Ridanpää et al., 2001), MUL is caused by mutations in the E3 ubiquitin ligase TRIM37 (Avela et al., 2000; Kallijärvi et al., 2005). Both CHH and MUL patients have increased cancer risk, and expression of *RMRP* and TRIM37 are implicated in cancer progression (Hao and Zhou, 2022; Przanowski et al., 2020).

*RMRP* dysfunction affects the expression of cell cycle regulating genes (Vakkilainen et al., 2019; Gao et al., 2025; Sun et al., 2019; Thiel et al., 2005), p53 protein level (Chen et al., 2021), and ribosome synthesis (Robertson et al., 2022), but how pathogenic variants of *RMRP* cause proliferative defects and increased cancer risk remains unclear. Despite reports on increased duration of the late cell cycle (Vakkilainen et al., 2019; Gill et al., 2004), a characterization of functional cellular states based on protein levels and signaling status is lacking.

Analysis of signaling trajectories requires multiplexed information from single cells. Due to the large number of available antibodies, immunofluorescence can detect a wide selection of proteins and post-translational modifications. In addition to the advantage of subcellular resolution, we have previously showed that immunofluorescence can be quantitative and be used to estimate temporal signaling dynamics (Akopyan et al., 2014). However, quantitative immunofluorescence is restricted by the number of different antibodies that can be used on one cell.

Different approaches have been used to allow detection of more than one staining in fluorescence microscopy. A combination of antibodies from different species, followed by secondary antibodies coupled to different fluorophores, is routinely used. More recently, sequential or cyclical imaging has been established (Hickey et al., 2022). These methods rely on blocking the signal after imaging to allow another round of staining and imaging. Various methods have been established, including bleaching fluorophores (Schubert et al., 2006; Radtke et al., 2020), labeling antibodies with DNA barcodes (Goltsev et al., 2018; Saka et al., 2019) and elution of bound antibodies (Lan et al., 1995; Pirici et al., 2009). Methods that depend on bleaching and DNA barcodes require labelling of primary antibodies, excluding use of off-the-shelf antibodies and amplification by secondary antibodies. Antibody elution balances efficient antibody removal with loss of epitopes (Klevanski et al., 2020), and requires strategies to minimize imaging-induced cross-linking of antibodies (Gut et al., 2018).

We have developed cumulative microscopy, a method that relies on sequential imaging which retains the signal of previous stainings and therefore does not require bleaching, quenching or elution steps. We find that cumulative microscopy is quantitative and can support neural network-based analysis as well as identification of cellular states and trajectories between states. Using cumulative microscopy on fibroblasts from patients with rare genetic disorders, we find that a subset of fibroblasts from CHH patients show spontaneous replication stress. We note that replication stress potentially could explain both proliferation defects and cancer prevalence in CHH patients.

## Results and Discussion

We developed cumulative microscopy, a method to multiplex fluorescence microscopy by computationally extracting individual signals from sequential imaging. During consecutive imaging cycles, different primary antibodies from the same species are used with the same secondary antibody. The resulting images show an additive staining over each imaging cycle. To retrieve the signal of one specific antibody, the image of the previous imaging cycle is subtracted (Figure 1A). To control for bleaching and additive binding of secondary antibodies, the subtraction is calibrated by control images in which the last primary antibody is omitted (Supplementary figure 1). Due to the calibration step, exposure conditions can be changed between imaging cycles, avoiding eventual saturation of the signal.

**Figure 1:**
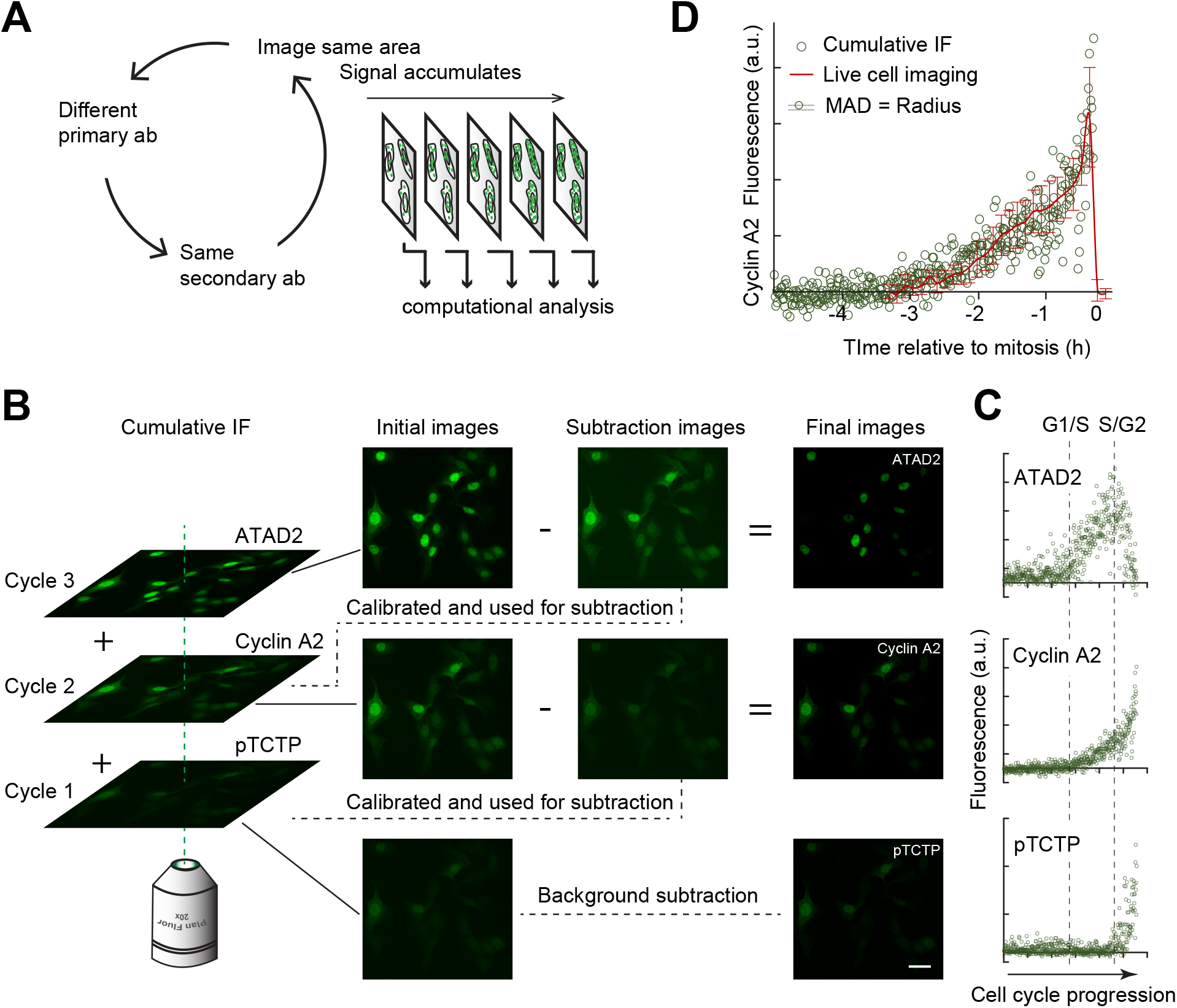
Cumulative microscopy. A, schematic of cumulative microscopy. B, workflow for cumulative microscopy. Images to the right show resulting subtractions after cumulative staining with ATAD2, Cyclin A2, and pTCTP antibodies. Scale bar 40 μm. C, quantification after imaging data as in B, with the difference that estimation of scaling function and subtraction is performed on measurements of nuclei rather than on full images. Graphs show measurements for individual cells sorted *in silico* for cell cycle progression. D, comparison of Cyclin A2 levels based on live-cell imaging of Cyclin A2-YFP gene-targeted RPE cells (red line) and time-estimated Cyclin A2 levels in RPE cells based on data from C (green circles). Size of circles indicate estimated technical variability (mean absolute deviation, MAD).

The subtractions in consecutive imaging cycles can either be performed directly on the images or on measurements from the images (Figure 1A, B, and supplementary figure 2). From a practical perspective, we find that the latter more easily can correct for a shift in position that may occur if the same specimen is repeatedly replaced in a microscope, and therefore allows processing without image registration (Supplementary figure 2).

We developed a quantitative estimate of the uncertainty introduced by cumulative microscopy that is readily accessible from the calibration of each imaging cycle (Supplementary figure 3). We note that the uncertainty can be small compared to variation between cells as assessed both by live cell imaging and quantitative immunofluorescence (Figure 1C, D, and supplementary figure 4). In agreement, cumulative microscopy showed a similar expression pattern over time of a cell cycle regulated protein as one round of immunofluorescence, while allowing separation of distinct expression patterns of other proteins (Figure 1C and supplementary figure 4). Similarly, both expression pattern over time and variation between cells are similar between cumulative microscopy for Cyclin A2 and live-cell imaging of Cyclin A2-YFP (Figure 1D). Taken together, this indicates that cumulative microscopy can be used to quantitatively assess antibody stainings. For a more detailed discussion of the quantitative estimate of variability and possible error sources, please see supplementary text.

We used cumulative microscopy on a collection of primary fibroblasts from individuals with monogenic growth disorders with cell proliferative defects and from matched controls (Supplementary figure 5A). We chose the antibodies to detect key proteins regulating cell cycle progression and stress response (Supplementary figure 5B). At the same time, we recorded DAPI intensity to estimate DNA content, as well as nuclear area and EdU incorporation the last hour before fixation. After quantification of the signals from each cell, we trained a neural network to detect different cell cycle phases. Not surprisingly considering the measurements of DNA content, EdU incorporation and main cell cycle regulators, the neural network could accurately identify cell cycle phases (Figure 2A-C). This shows that data from cumulative microscopy can be used for neural-network-based predictions.

**Figure 2:**
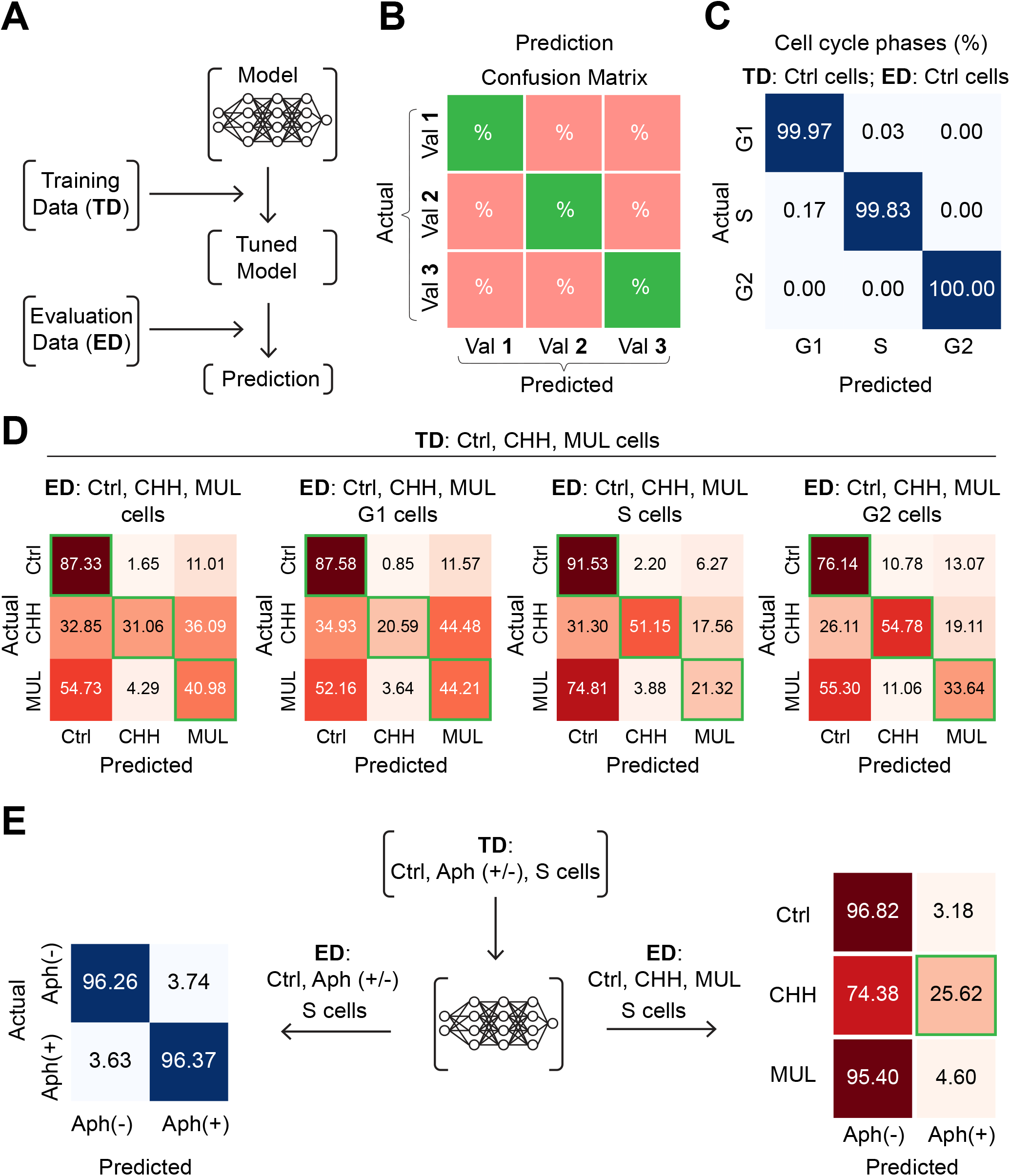
Neural network-based analysis of cumulative microscopy data. A, schematic of analysis. B, schematic of confusion matrix for visualization of results. C, model learning and prediction of cell cycle phase based on cumulative microscopy data on untreated control cells. D, model learning and prediction of cell type based on cumulative microscopy data on untreated control, CHH, and MUL cells. Left matrix shows prediction for all cells, remaining matrices show prediction separated for individual cell cycle phases. E, left matrix shows learning and prediction based on control S phase cells treated with or without aphidicolin. Right matrix shows learning based on control S phase cells treated with or without aphidicolin, and prediction using untreated control, CHH, and MUL S phase cells.

We next sought to test whether a neural network based on cell cycle and stress response markers could identify cells from different disorders. We therefore trained the network on data from primary fibroblasts from CHH and MUL patients and matched controls. Interestingly, although the data is based solely on cell cycle and stress response markers, 31 and 41% of CHH and MUL cells were correctly identified (Figure 2D). Whereas identification of MUL cells was most efficient in G1 phase, identification of CHH cells was most efficient in S and G2 phases (Figure 2D). This indicates that at least a subset of CHH and MUL cells have stress and cell cycle characteristics that separate them from control cells and from each other.

As CHH cells, but not MUL cells, were mainly identified in S and G2 phases, we reasoned that these cells could exhibit some difference during DNA replication. We therefore perturbed DNA replication by adding a low level of the DNA-polymerase inhibitor Aphidicolin. After cumulative microscopy, we trained a neural network to score whether S phase control fibroblasts were exposed to aphidicolin (Figure 2E, left). We next used the network trained on control cells treated or not with aphidicolin to score CHH and MUL cells in the absence of aphidicolin.

Interestingly, although CHH cells were not treated with aphidicolin, the neural network scored 26% of S phase cells as treated with aphidicolin. In contrast, control and MUL cells were not identified as treated with aphidicolin (Figure 2E, right). This suggests that at least a subset of S phase CHH cells, but not MUL cells, are sharing properties with control cells treated with Aphidicolin.

To visualize the cellular states, we performed a uniform manifold approximation and projection (UMAP) on all data combined and noted that fibroblasts from different individuals aligned in similar patterns (Figure 3A). These patterns resembled distinct cellular states depending on cell cycle and stress responses. Importantly, by overlaying the individual antibody patterns, it was possible to identify states as replication stress, DNA damage checkpoints, and cell cycle exit (Figure 3B-E). We identified the predominant trajectories of the unperturbed cell cycle in untreated cells (blue arrows), and the predominant trajectories in cells treated with a low level of aphidicolin (red arrows) (Figure 3F). As expected, we find that S phase cells gain a separate trajectory after Aphidicolin addition, characterized by reduced EdU incorporation and increased γH2AX levels, consistent with cells experiencing impeded DNA replication. In addition, we note that both G1 cells and G2 cells gained separate trajectories, characterized among others by expression of p53 and p21.

**Figure 3:**
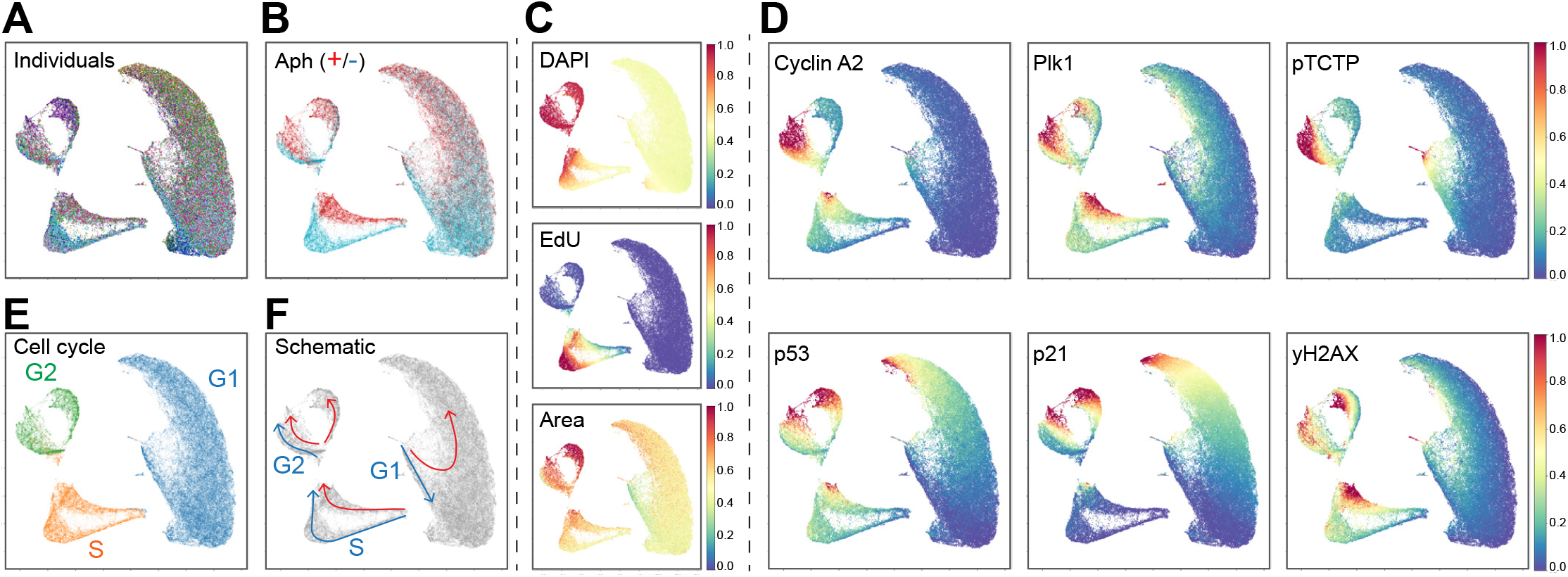
Assessing cell states after cumulative microscopy. A, UMAP of cumulative microscopy data of in total 96120 primary cells from 15 individuals, with or without 48h exposure to aphidicolin. Colors indicate the 30 different conditions. B, UMAP from A, indicating with or without exposure to aphidicolin. C, UMAP from A, indicating DAPI, EdU levels, and nuclear area. D, UMAP from A, indicating the antibodies used for cumulative microscopy. E, UMAP from A, indicating cell cycle stage based on DAPI and EdU data. F, UMAP from A, arrows indicate assessed main cell cycle trajectories. Blue indicates unperturbed cells, and red indicates after aphidicolin exposure.

We next gated the distributions of cells from each individual in the UMAP. We noted that cells from 4 individuals diagnosed with CHH showed a different pattern compared to control cells (Figure 4A). First, an increased fraction of the untreated CHH cells colocalized with replicating cells treated with the DNA polymerase inhibitor aphidicolin (Figure 3B and 4B). Similar to cells treated with aphidicolin, these CHH cells showed an accumulation of γH2AX, indicating DNA damage, and increased Cyclin A2 and P1k1 levels compared to their position in S phase, indicating that the cell cycle engine had proceeded further than the replication of DNA (Figure 4B and supplementary figure 6) (Lemmens et al., 2018). Similarly, CHH cells in S phase showed a reduction of EdU incorporation, although to a lesser extent compared to cells treated with aphidicolin that directly inhibits DNA polymerase (Figure 4C and supplementary figure 6). Thus, a subset of CHH cells experiences replication stress.

**Figure 4:**
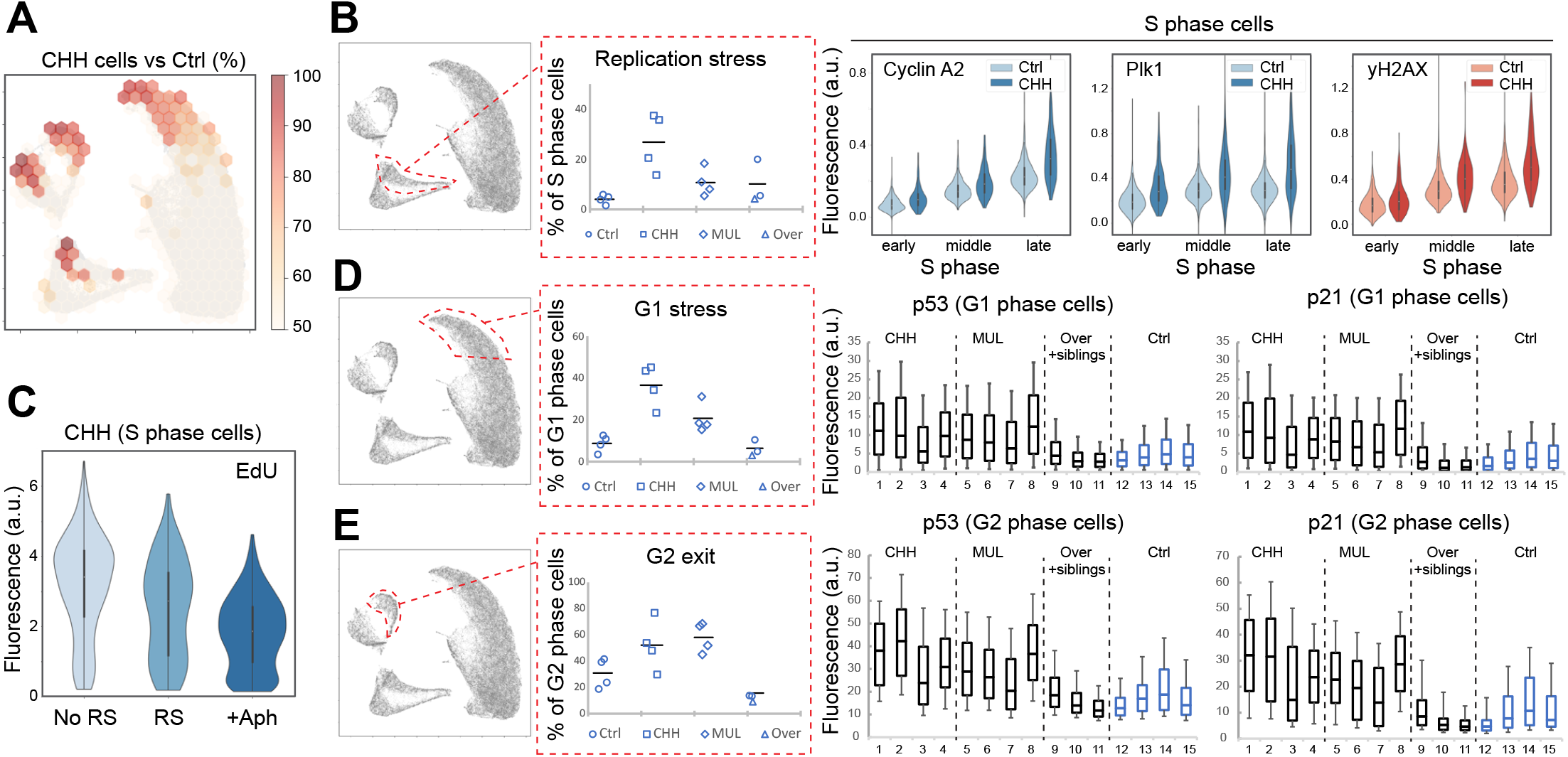
A subset of CHH cells show replication stress and cell cycle exit in both G1 and G2 phases. A, accumulation of CHH cells. Graph shows percentage of CHH cells versus Ctrl cells. The number of cells was normalized and binned into hexagons through the UMAP. B, left, gate in UMAP to depict replication stress. Right, distribution of indicated measurements in S phase cells. C, EdU incorporation in S phase for different fractions of CHH cells: No RS - cells colocalized with untreated Ctrl replicating cells, RS - cells colocalized with cells treated with aphidicolin and fraction (+Aph) - cells treated with aphidicolin. D, left, gate in UMAP to depict G1 stress. Right, distribution of indicated measurements in G1 phase cells. See supplementary figure 8 for P values. E, left, gate in UMAP to depict cell cycle exit from G2 phase. Right, distribution of indicated measurements in G2 phase cells. See supplementary figure 8 for P values.

Replication stress can lead to p53- and p21-dependent cell cycle exit in the following G2 phase or, if the cells pass through mitosis, to a p53 and p21-dependent cell cycle block in the following G1 phase (Hornsveld et al., 2021; Arora et al., 2017). We found that CHH cells accumulated in the corresponding regions in the UMAP and that both G1 and G2 cells contained increased levels of p53 and p21 (Figure 4D, E). Similar to CHH cells, we found that cells from 4 individuals diagnosed with Mulibrey nanism contained upregulated p53 and p21 and accumulated in the regions for G1 cell cycle arrest and G2 cell cycle exit (Figure 3C, D). However, in contrast to CHH, MUL cells did not show increased colocalization with aphidicolin treated cells (Figure 4A). This difference possibly reflects underlying mutations for the conditions, as MUL cells contain mutations in *TRIM37* that is involved in ensuring bipolar spindle assembly in mitosis (Balestra et al., 2021; Avela et al., 2000; Bellaart et al., 2025; Yeow et al., 2025). Cells from an individual with an unspecified overgrowth syndrome did not accumulate in regions for cell cycle exit or arrest, suggesting faithful execution of the cell cycle. We conclude that CHH and MUL cells both show a stress-induced block to proliferation, but by different mechanisms.

To independently test the results obtained by cumulative microscopy, we monitored the cells using time-lapse microscopy. Whereas both MUL and CHH cells showed increased number of cells that are not dividing, only CHH cells showed an increased cell cycle duration, in agreement with spontaneous replication stress followed by checkpoint activation in G2 phase (Vakkilainen et al., 2019) (Supplementary figures 5, 7).

Replication stress has emerged as a key driver of genome instability in cancer, and a common feature of multiple oncogenes (Halazonetis et al., 2008; Gaillard et al., 2015). More recently, treatment options that target replication stress have been considered (Cybulla and Vindigni, 2023). We note that replication stress potentially could underlie both proliferative defects and increased cancer risk in CHH patients.

CHH patients carry biallelic pathogenic variants in the *RMRP* gene, but whether and how *RMRP* contributes to replication stress requires further investigation. However, we note that an inverse correlation exists between *RMRP* expression and a replication stress score in established cell lines (Supplementary figure 9).

We conclude that cumulative microscopy is a straightforward method to multiplex immunofluorescence that does not require elution or bleaching steps. The method is quantitative, can be performed using microscopes that are commonly available, does not require antibody conjugation steps, and can be used to support multi-labeling-based identification of cellular states. Importantly, although tested for antibody-based labelling here, cumulative microscopy can in theory be expanded to other forms of microscopy that involve quantification of a fluorescent signal.

## Materials and Methods

### Primary fibroblasts

The collection of patient and control skin biopsies for research purposes was approved by the Institutional Research Ethics Committee of the Helsinki University Hospital, Finland (HUS/836/2018, HUS/1190/2018). All the participants gave an informed consent.

Forearm skin biopsies were harvested from the mutation-positive subjects (four with CHH and four with MUL) and from four age- and sex matched controls. Further, skin biopsies were harvested from a subject with an unspecified overgrowth syndrome, as well as from two siblings without overgrowth. Harvesting of skin biopsies and isolation of fibroblasts was performed as described in (Pekkinen et al., 2019).

### Cell culture

RPE and RPE - CyclinA2-YFP cells (Silva Cascales et al., 2021) were cultured in DMEM-F12 medium plus GlutaMAX (Invitrogen) supplemented with 10% FBS (HyClone) and 1% Penicillin-Streptomycin (HyClone) and primary human fibroblasts were cultured in DMEM medium plus GlutaMAX (Invitrogen) supplemented with 10% FBS (HyClone) and 1% Penicillin-Streptomycin (HyClone). Cells were maintained at 37^0^C and 5 % CO_2_.

Before experiments cells were seeded in 96-well imaging plates for overnight.

*Reagents*
| Reagent | Source | Identifier |
| --- | --- | --- |
| Ascorbic acid | Sigma Aldrich | A4403 |
| Aphidicolin | Sigma Aldrich | A4487 |
| Azid Fluorophore (Alexa 647) | Invitrogen | A10277 |
| BSA | Sigma Aldrich | A4503 |
| CuSO<sub>4</sub> | Sigma Aldrich | 451657 |
| DAPI | ThermoFisher | 62248 |
| DMSO | Sigma Aldrich | D4540 |
| EdU (5-ethynyl-2'-deoxyuridine) | ThermoFisher | E10187 |
| Goat anti-Mouse conjugated to Alexa Fluor™ 568 | Invitrogen | A11004 |
| Goat anti-Rabbit conjugated to Alexa Fluor™ 488 | Invitrogen | A11034 |
| Methanol | Sigma Aldrich | 32213N |
| Mouse anti p53 | Santa Cruz | sc-126 |
| Mouse anti Plk1 | Santa Cruz | sc-17783 |
| Mouse anti gH2AX | Millipore | 05-636 |
| PBS | Gibco | 14190-144 |
| Rabbit anti ATAD2 | Sigma Aldrich | HPA029424 |
| Rabbit anti Cyclin A2 | Atlas antibodies | HPA020626 |
| Rabbit anti p21 | Cell Signaling | 2947 |
| Rabbit anti Phospho-TCTP | Cell Signaling | 5251 |
| TBS (Tris buffered saline pH 7.6) | ThermoFisher | J62662 |
| TBS-T (TBS containing 0.1% Tween-20) | Sigma Aldrich | P1379 |
| Tris (pH 6.8) | Sigma Aldrich | T1503 |

### Reagents

#### Immunofluorescence and microscopy

Immunofluorescence was performed as in (Akopyan et al., 2016). An ImageXpress microscopy system equipped with a CoolSNAP HQ camera (Molecular Devices/LLC, Sunnyvale, CA, USA) and a Nikon 20× Plan Fluor ELWD NA 0.45 objective was used to acquire fluorescent images.

#### Live-Cell Imaging

Primary fibroblasts were followed on a Satorius IncuCyte S3 Long-term imaging inside a cell culture incubator system using a 20× NA 0.45 air objective for 60 hours. Images were acquired every 1 hour.

RPE - CyclinA2-YFP cells were followed on a Leica DMI6000 Imaging System using a 20× NA 0.40 air objective. Images were acquired every 30 min. The medium of the cells was changed 24 hours prior to imaging to CO_2_-independent medium Leibovitz L-15 (ThermoFisher) supplemented with 6% or 10% heat-inactivated FBS and 1% Penicillin-Streptomycin (HyClone).

#### Image processing

Images were processed using ImageJ (http://rsb.info.nih.gov/ij/). Background subtraction was performed using background functions as in (Akopyan et al., 2016). Nuclei were segmented based on DAPI signal and integrated intensities of the nuclei were quantified. Pairwise registration of nuclei from two imaging cycles was performed by a custom-written script (ImageJ) that compared x-y position and nuclear area. Linear regression and pairwise measurement were performed in Python using *scipy*.*stats*.*linregress*.

Time estimation from fixed cells was performed by ordering cells based on increasing level of Cyclin A2 and DAPI as in (Akopyan et al., 2014).

#### Dimensionality reduction algorithm and simulations

Uniform Manifold Approximation and Projection (UMAP) and visualization were performed in Python using *sklearn*.*datasets, sklearn*.*model_selection, sklearn*.*preprocessing, matplotlib*.*pyplot, numpy, seaborn, pandas*, and *colorcet*.

Simulations for relation of R-square value to normalized standard deviation were performed in Python using: *matplotlib*.*pyplot, numpy, scipy*.*stats, seaborn, pandas, pathlib*, and *math*.

#### Neural network architecture and training procedure

A feedforward neural network was implemented using PyTorch (version 2.5.1) to classify cell types, cell phases or conditions based on input data. The model architecture consists of an input layer with 9 neurons, followed by three fully connected hidden layers with 8, 9, and 9 neurons respectively. Each hidden layer uses the Rectified Linear Unit (ReLU) activation function to introduce non-linearity. The final output layer comprises 2 neurons corresponding to the binary classification task (e.g., Aphidicolin-treated vs. not treated cells) or 3 neurons in case of classifying cell types or cell phases.

Prior to training, the dataset was split into training and test sets using an 80:20 ratio with a fixed random seed for reproducibility. Input features were standardized using the StandardScaler from the scikit-learn library to ensure consistent scaling across all features. The model was trained using the Adam optimizer with a learning rate of 0.01. The loss function used was cross-entropy loss, suitable for multi-class classification tasks. Training was conducted over 600 epochs, and the model’s performance was monitored by tracking the training loss at regular intervals.

The model was trained and evaluated on separate datasets to assess its generalizability across different experimental conditions.

#### Replication Stress Score

The replication stress (Repstress) score was calculated by summing the expression levels of 17 signature genes, each weighted according to previously published coefficients (Takahashi et al., 2022). Gene expression values were Z score-normalized across all genes within each cell line. The resulting Repstress scores were then Z score-normalized across all samples to enable cross-sample comparison.

RNA-seq data for 675 commonly used human cancer cell lines were downloaded from the EMBL-EBI Expression Atlas (https://www.ebi.ac.uk/gxa/experiments/E-MTAB-2706/Downloads). Only cell lines with non-zero expression values for RMRP, TRIM37, and all 17 signature genes were included in the analysis, resulting in a final dataset of 180 cell lines spanning 49 cancer types.

## Funding

The Swedish Research Council 2019-01594 (AL)

The Swedish Cancer Society 2021/1742 (AL)

The Swedish Childhood Cancer Foundation PR2021-0090 (OM, AL)

Stockholm’s County Council (ALF) 502492 (OM)

The Swedish Research Council 2022-00800 (OM)

Folkhälsan Research Foundation 101009 (OM)

Finska Läkaresällskapet (SK, OM)

Stiftelsen Dorothea Olivia, Karl Walter och Jarl Walter Perkléns minne (SK)

Sällsyntafonden (FT)

## Author contributions

KA and AL conceived the method. KA established the method, performed the imaging experiments and analyzed the data. SK, SV, FT, and MP recruited the patients and established the primary fibroblasts. OM and AL supervised. KA and AL wrote the manuscript. All authors revised the manuscript.

## Conflict of interests

The authors declare that they have no conflict of interests.

## Code availability

Example code to perform cumulative microscopy is available at https://github.com/KAkopyan/CumulativeMicroscopy

## References

Akopyan, K., A. Lindqvist, and E. Müllers. 2016. Cell Cycle Dynamics of Proteins and Post-translational Modifications Using Quantitative Immunofluorescence. Methods Mol Biol. 1342:173–183. doi:10.1007/978-1-4939-2957-39.

Akopyan, K., H. Silva Cascales, E. Hukasova, A.T. Saurin, E. Müllers, H. Jaiswal, D.A.A. Hollman, G.J.P.L. Kops, R.H. Medema, and A. Lindqvist. 2014. Assessing kinetics from fixed cells reveals activation of the mitotic entry network at the S/G2 transition. Mol Cell. 53:843–853. doi:10.1016/j.molcel.2014.01.031.

Arora, M., J. Moser, H. Phadke, A.A. Basha, and S.L. Spencer. 2017. Endogenous Replication Stress in Mother Cells Leads to Quiescence of Daughter Cells. Cell Reports. 19:1351–1364. doi:10.1016/j.celrep.2017.04.055.

Avela, K., M. Lipsanen-Nyman, N. Idänheimo, E. Seemanová, S. Rosengren, T.P. Mäkelä, J. Perheentupa, A.D. Chapelle, and A.E. Lehesjoki. 2000. Gene encoding a new RING-B-box-Coiled-coil protein is mutated in mulibrey nanism. Nat Genet. 25:298–301. doi:10.1038/77053.

Balestra, F.R., A. Domínguez-Calvo, B. Wolf, C. Busso, A. Buff, T. Averink, M. Lipsanen-Nyman, P. Huertas, R.M. Ríos, and P. Gönczy. 2021. TRIM37 prevents formation of centriolar protein assemblies by regulating Centrobin. Elife. 10:e62640. doi:10.7554/eLife.62640.

Bellaart, A., A. Brambila, J. Xu, F. Mendez Diaz, A. Deep, J. Anzola, F. Meitinger, M. Ohta, K.D. Corbett, A. Desai, and K. Oegema. 2025. TRIM37 prevents ectopic spindle pole assembly by peptide motif recognition and substrate-dependent oligomerization. Nat Struct Mol Biol. doi:10.1038/s41594-025-01562-0.

Chen, Y., Q. Hao, S. Wang, M. Cao, Y. Huang, X. Weng, J. Wang, Z. Zhang, X. He, H. Lu, and X. Zhou. 2021. Inactivation of the tumor suppressor p53 by long noncoding RNA RMRP. Proc Natl Acad Sci U S A. 118:e2026813118. doi:10.1073/pnas.2026813118.

Cybulla, E., and A. Vindigni. 2023. Leveraging the replication stress response to optimize cancer therapy. Nat Rev Cancer. 23:6–24. doi:10.1038/s41568-022-00518-6.

Gaillard, H., T. García-Muse, and A. Aguilera. 2015. Replication stress and cancer. Nat Rev Cancer. 15:276-289. doi:10.1038/nrc3916.

Gao, J., J. Zheng, S. Chen, S. Lin, and S. Duan. 2025. RMRP variants inhibit the cell cycle checkpoints pathway in cartilage-hair hypoplasia. Mol Med Rep. 31:81. doi:10.3892/mmr.2025.13446.

Gill, T., T. Cai, J. Aulds, S. Wierzbicki, and M.E. Schmitt. 2004. RNase MRP cleaves the CLB2 mRNA to promote cell cycle progression: novel method of mRNA degradation. Mol Cell Biol. 24:945–953. doi:10.1128/MCB.24.3.945-953.2004.

Goltsev, Y., N. Samusik, J. Kennedy-Darling, S. Bhate, M. Hale, G. Vazquez, S. Black, and G.P. Nolan. 2018. Deep Profiling of Mouse Splenic Architecture with CODEX Multiplexed Imaging. Cell. 174:968–981.e15. doi:10.1016/j.cell.2018.07.010.

Gut, G., M.D. Herrmann, and L. Pelkmans. 2018. Multiplexed protein maps link subcellular organization to cellular states. Science. 361:eaar7042. doi:10.1126/science.aar7042.

Haendel, M., N. Vasilevsky, D. Unni, C. Bologa, N. Harris, H. Rehm, A. Hamosh, G. Baynam, T. Groza, J. McMurry, H. Dawkins, A. Rath, C. Thaxon, G. Bocci, M.P. Joachimiak, S. Köhler, P.N. Robinson, C. Mungall, and T.I. Oprea. 2020. How many rare diseases are there? Nat Rev Drug Discov. 19:77–78. doi:10.1038/d41573-019-00180-y.

Halazonetis, T.D., V.G. Gorgoulis, and J. Bartek. 2008. An oncogene-induced DNA damage model for cancer development. Science. 319:1352–1355. doi:10.1126/science.1140735.

Hao, Q., and X. Zhou. 2022. The emerging role of long noncoding RNA RMRP in cancer development and targeted therapy. Cancer Biol Med. 19:140–146. doi:10.20892/j.issn.2095-3941.2021.0577.

Hickey, J.W., E.K. Neumann, A.J. Radtke, J.M. Camarillo, R.T. Beuschel, A. Albanese, E. McDonough, J. Hatler, A.E. Wiblin, J. Fisher, J. Croteau, E.C. Small, A. Sood, R.M. Caprioli, R.M. Angelo, G.P. Nolan, K. Chung, S.M. Hewitt, R.N. Germain, J.M. Spraggins, E. Lundberg, M.P. Snyder, N.L. Kelleher, and S.K. Saka. 2022. Spatial mapping of protein composition and tissue organization: a primer for multiplexed antibody-based imaging. Nat Methods. 19:284–295. doi:10.1038/s41592-021-01316-y.

Hornsveld, M., F.M. Feringa, L. Krenning, J. van den Berg, L.M.M. Smits, N.B.T. Nguyen, M.J. Rodríguez-Colman, T.B. Dansen, R.H. Medema, and B.M.T. Burgering. 2021. A FOXO-dependent replication checkpoint restricts proliferation of damaged cells. Cell Rep. 34:108675. doi:10.1016/j.celrep.2020.108675.

Kallijärvi, J., U. Lahtinen, R.H ämäläinen, M. Lipsanen-Nyman, J.J. Palvimo, and A.-E. Lehesjoki. 2005. TRIM37 defective in mulibrey nanism is a novel RING finger ubiquitin E3 ligase. Exp Cell Res. 308:146–155. doi:10.1016/j.yexcr.2005.04.001.

Klevanski, M., F. Herrmannsdoerfer, S. Sass, V. Venkataramani, M. Heilemann, and T. Kuner. 2020. Automated highly multiplexed super-resolution imaging of protein nano-architecture in cells and tissues. Nat Commun. 11:1552. doi:10.1038/s41467-020-15362-1.

Lan, H.Y., W. Mu, D.J. Nikolic-Paterson, and R.C. Atkins. 1995. A novel, simple, reliable, and sensitive method for multiple immunoenzyme staining: use of microwave oven heating to block antibody crossreactivity and retrieve antigens. J Histochem Cytochem. 43:97–102. doi:10.1177/43.1.7822770.

Lemmens, B., N. Hegarat, K. Akopyan, J. Sala-Gaston, J. Bartek, H. Hochegger, and A. Lindqvist. 2018. DNA Replication Determines Timing of Mitosis by Restricting CDKl and PLKl Activation. Mol Cell. 71:117–128.e3. doi:10.1016/j.molcel.2018.05.026.

Movasat, H., E. Giacopino, A. Shahdoost, Y.D. Nokoorani, A.H. Abrbekouh, Y. Tahamtani, and N. Shakiba. 2025. A systems view of cellular heterogeneity: Unlocking the “wheel of fate.” cels. 0. doi:10.1016/j.cels.2025.101300.

Pekkinen, M., P.A. Terhal, L.D. Botto, P. Henning, R.E. Mäkitie P. Roschger, A. Jain, M. Kol, M.A. Kjellberg, E.P. Paschalis, K. van Gassen, M. Murray, P. Bayrak-Toydemir, M.K. Magnusson, J. Jans, M. Kausar, J.C. Carey, P. Somerharju, U.H. Lerner, V.M. Olkkonen, K. Klaushofer, J.C. Holthuis, and O.M äkitie. 2019. Osteoporosis and skeletal dysplasia caused by pathogenic variants in SGMS2. JCI Insight. 4:e126180. 126180. doi:10.1172/jci.insight.126180.

Pirici, D., L. Mogoanta, S. Kumar-Singh, I. Pirici, C. Margaritescu, C. Simionescu, and R. Stanescu. 2009. Antibody Elution Method for Multiple Immunohistochemistry on Primary Antibodies Raised in the Same Species and of the Same Subtype. J Histochem Cytochem. 57:567–575. doi:10.1369/jhc.2009.953240.

Przanowski, P., S. Lou, R.D. Tihagam, T. Mondal, C. Conlan, G. Shivange, I. Saltani, C. Singh, K. Xing, B.B. Morris, M.W. Mayo, L. Teixeira, J. Lehmann-Che, J. Tushir-Singh, and S. Bhatnagar. 2020. Oncogenic TRIM37 links chemoresistance and metastatic fate in triple-negative breast cancer. Cancer Res. 80:4791–4804. doi:10.1158/0008-5472.CAN-20-1459.

Radtke, A.J., E. Kandov, B. Lowekamp, E. Speranza, C.J. Chu, A. Gola, N. Thakur, R. Shih, L. Yao, Z.R. Yaniv, R.T. Beuschel, J. Kabat, J. Croteau, J. Davis, J.M. Hernandez, and R.N. Germain. 2020. IBEX: A versatile multiplex optical imaging approach for deep phenotyping and spatial analysis of cells in complex tissues. Proceedings of the National Academy of Sciences. 117:33455–33465. doi:10.1073/pnas.2018488117.

Rafelski, S.M., and J.A. Theriot. 2024. Establishing a conceptual framework for holistic cell states and state transitions. Cell. 187:2633–2651. doi:10.1016/j.cell.2024.04.035.

Ridanpää, M., H. van Eenennaam, K. Pelin, R. Chadwick, C. Johnson, B. Yuan, W. vanVenrooij, G. Pruijn, R. Salmela, S. Rockas, O.M äkitie, I. Kaitila, and A. de la Chapelle. 2001. Mutations in the RNA component of RNase MRP cause a pleiotropic human disease, cartilage-hair hypoplasia. Cell. 104:195–203. doi:10.1016/s0092-8674(01)00205-7.

Robertson, N., V. Shchepachev, D. Wright, T.W. Turowski, C. Spanos, A. Helwak, R. Zamoyska, and D. Tollervey. 2022. A disease-linked lncRNA mutation in RNase MRP inhibits ribosome synthesis. Nat Commun. 13:649. doi:10.1038/s41467-022-28295-8.

Saka, S.K., Y. Wang, J.Y. Kishi, A. Zhu, Y. Zeng, W. Xie, K. Kirli, C. Yapp, M. Cicconet, B.J. Beliveau, S.W. Lapan, S. Yin, M. Lin, E.S. Boyden, P.S. Kaeser, G. Pihan, G.M. Church, and P. Yin. 2019. Immuno-SABER enables highly multiplexed and amplified protein imaging in tissues. Nat Biotechnol. 37:1080–1090. doi:10.1038/s41587-019-0207-y.

Schubert, W., B. Bonnekoh, A.J. Pommer, L. Philipsen, R. Böckelmann, Y. Malykh, H. Gollnick, M. Friedenberger, M. Bode, and A.W.M. Dress. 2006. Analyzing proteome topology and function by automated multidimensional fluorescence microscopy. Nat Biotechnol. 24:1270–1278. doi:10.1038/nbt1250.

Silva Cascales, H., K. Burdova, A. Middleton, V. Kuzin, E. Müllers, H. Stoy, L. Baranello, L. Macurek, and A. Lindqvist. 2021. Cyclin A2 localises in the cytoplasm at the S/G2 transition to activate PLK1. Life Sci Alliance. 4:e202000980. doi:10.26508/1sa.202000980.

Sun, X., R. Zhang, M. Liu, H. Chen, L. Chen, F. Luo, D. Zhang, J. Huang, F. Li, Z. Ni, H. Qi, N. Su, M. Jin, J. Yang, Q. Tan, X. Du, B. Chen, H. Huang, S. Chen, L. Yin, X. Xu, C. Deng, L. Luo, Y. Xie, and L. Chen. 2019. Rmrp Mutation Disrupts Chondrogenesis and Bone Ossification in Zebrafish Model of Cartilage-Hair Hypoplasia via Enhanced Wnt/β-Catenin Signaling. J Bone Miner Res. 34:2101–2116. doi:10.1002/jbmr.3820.

Thiel, C.T., D. Horn, B. Zabel, A.B. Ekici, K. Salinas, E. Gebhart, F. Rüschendorf, H. Sticht, J. Spranger, D. Müller, C. Zweier, M.E. Schmitt, A. Reis, and A. Rauch. 2005. Severely incapacitating mutations in patients with extreme short stature identify RNA-processing endoribonuclease RMRP as an essential cell growth regulator. Am J Hum Genet. 77:795–806. doi:10.1086/497708.

Vakkilainen, S., T. Skoog, E. Einarsdottir, A. Middleton, M. Pekkinen, T. Öhman, S. Katayama, K. Krjutškov, P.E. Kovanen, M. Varjosalo, A. Lindqvist, J. Kere, and O. Mäkitie. 2019. The human long non-coding RNA gene RMRP has pleiotropic effects and regulates cell-cycle progression at G2. Sci Rep. 9:13758. doi:10.1038/s41598-019-50334-6.

Yeow, Z.Y., S. Sarju, F.-C. Chang, L.Y. Xu, M. van Breugel, and A.J. Holland. 2025. Mesoscale regulation of microtubule-organizing centers by the E3 ligase TRIM37. Nat Struct Mol Biol. doi:10.1038/s41594-025-01540-6.

